# Role of pioneer neurons and neuroblast behaviors on otic ganglion assembly

**DOI:** 10.1101/2023.03.30.534903

**Authors:** A Bañón, B Alsina

## Abstract

Cranial ganglia are aggregates of sensory neurons that mediate distinct types of sensation. It is little understood how individual neurons coalesce, distribute and shape the ganglion. The statoacoustic ganglion (SAG) displays several lobes spatially arranged to properly connect with hair cells of the inner ear. To investigate the cellular behaviors involved in the 3D organization of the SAG, we use high resolution confocal imaging of single cell labeled zebrafish neuroblasts (NB), photoconversion, photoablation and genetic perturbations. We find that otic NB delaminate out of the otic epithelium in an EMT-like manner, rearranging apical polarity and primary cilia proteins. We also show that, once delaminated, NB migrate directionally and actively, requiring RhoGTPases. Interestingly, cell tracking of individual delaminated NB reveals that NB migrate and coalesce around a small population of pioneer SAG neurons. These pioneer SAG neurons are not from otic placode origin and populate the coalescence region before otic neurogenesis begins. Upon ablation of these cells, migratory pathways of delaminated NB are disrupted and, consequently, SAG shape is affected. Altogether, this work shows for the first time the role of pioneer SAG neurons in orchestrating SAG development.

**Summary Statement:** Little is known how cranial sensory ganglia organize in 3D. We unveil the repertoire of cellular behaviours underlying statoacoustic morphogenesis and its dependence on relevant pioneer neurons.

## Introduction

The inner ear is responsible for the senses of hearing and balance and is organized in two main structures, the epithelial labyrinth containing the hair cells and the statoacoustic ganglion (SAG), the inner ear cranial sensory ganglion. Through development, cranial sensory ganglia undergo cellular rearrangements in a process of morphogenesis to acquire its final functional shape. The SAG generates a particular architecture with three distinct lobes; two anterior lobes elongated dorsoventrally and a posterior lobe elongated anteroposteriorly that establish topographic projections with the sensory patches of the inner ear (Taberner et al., 2018). Thus, for a correct circuitry between hair cells and otic neurons, SAG morphogenesis must be tightly coordinated in time and space with otic tissue development.

Several studies have addressed the morphogenetic mechanisms underlying the 3D sculpturing of the inner ear labyrinth. In particular, the zebrafish has been an excellent model to live-image inner ear morphogenesis. By light sheet microscopy, it has been shown that zebrafish semicircular canal formation depends on ECM expansion and filopodia (Jones et al., 2022). On the other hand, lumen expansion requires anisotropic epithelial thinning and mechanical pulling of apical membranes and endolymphatic duct growth depends on lamellar projections (Hoijman et al., 2015; Swinburne et al., 2018). In contrast, live imaging of SAG neurons is a challenging task because neurons position behind the otic vesicle and rapidly compact into a ganglion. For that reason, SAG neurons have mainly been treated as a ganglionic unit and imaging and analysis of individual neuronal dynamics during SAG development has poorly been addressed.

The inner ear primordium, the otic placode, consists of an oval epithelium with an internal cavity or lumen. Otic progenitors display interkinetic nuclear migration and its apical membrane is oriented to the lumen and the basal membrane lines with the basal lamina (Alvarez et al., 1989). At the anteroventral quadrant of the otic vesicle the neurogenic domain emerges with the competence to specify neuronal precursors (Abelló et al., 2010; Hoijman et al., 2017). Neurogenesis initiates anteroventrally but progressively extends to posteromedial positions as observed by the temporal pattern of expression of *neurogenin1* (*neurog1*) in zebrafish, mouse and chick (Abelló et al., 2007; Bok et al., 2007, 2011; Radosevic et al., 2011). Later, Neurog1^+^ cells transit into Neurod^+^ cells and begin their delamination out of the otic epithelium (Ma et al., 1998; Sommer et al., 1996). In chick and zebrafish, it has been shown that the most anterolateral portion of the neurogenic domain generates vestibular neurons, whereas the posteromedial region gives rise to auditory neurons (Bell et al., 2008; Dyballa et al., 2017). Whether otic NB delamination follows an EMT program has been under debate. Some authors have suggested that core- EMT transcription factors and RhoGTPases (specially RhoB) are not used in this process of sensory neuron delamination in cranial placodes (Graham et al., 2007). Additionally, migratory capacity of cranial placode NB seems diminished compared to neural crest (Schlosser, 2008). Other evidences suggest the contrary, since *snail1b*, *cadherin6* and Sox9/10 family genes are expressed in delaminating Neurod^+^ otic NB and/or otic domains in zebrafish (Chiang et al., 2001; Dutton et al., 2009; Piloto & Schilling, 2010; Thisse et al., 1995).

In the developing central nervous system, sparse single cell labelling of membranes, nuclei and/or specific cellular components has revealed novel information on cell shape changes and behaviours during cell delamination, division or migration (Hadjivasiliou et al., 2019; Kasioulis et al., 2022; Moore et al., 2013). Radial glial cell migration in the cortical plate is a long-known mechanism by which newborn neurons use progenitor cell projections to properly migrate at later stages (Rakic, 1971a, 1971b; Shoukimas & Hinds, 1978). Specially studied in invertebrates is the modulation of neuronal dynamics during development by pioneer cells. These pioneer cells are defined as early born cells that have a scaffolding capacity to organize the tissue and preform the final configuration (Aigouy et al., 2008; Karkali et al., 2023; Wanner & Prince, 2013; Whitlock & Westerfield, 1998). Thus, a central question is how otic ganglion NB delaminate, migrate and coalesce (Dyballa et al., 2017; Hoijman et al., 2017) and whether other cells participate in SAG formation.

Here, we have addressed SAG morphogenesis by combining single cell labeling, photoconversion, photoablation and genetic perturbations. Our data suggest that indeed NB delamination is an EMT event. We could track individual NB during their migration before coalescence and revealed that is an active mechanism. We also find that otic NB migration depends on RhoGTPases. Interestingly, NB migrate and coalesce in a precise region populated by SAG pioneer neurons that arrived before neurogenic delamination in the otic placode begins. These pioneer SAG neurons recruit otic NB and are required for the organization of the SAG shape and size. In overall, the study provides novel data on how the SAG acquires its 3D organization and the underlying complex cellular behaviours of NB responsible for SAG development.

## Results

### Live imaging of otic neuroblasts during delamination reveal complex and dynamic cellular behaviors

Zebrafish NB delaminate out from the neurogenic domain depicted in red in an otic vesicle of 20 hpf (Fig. 1A) and generate the trilobular SAG behind the otic vesicle as visualized with the transgenic line *Tg(neurod:eGFP)* (Fig. 1A, green and Fig. 1B, right panel). In the ventral neurogenic domain, predelaminating NB remodel and acquire a different shape than cells in the non-neurogenic domain, which are more columnar (compare green with purple pseudocolored cells Fig. 1B).

**Figure 1.**
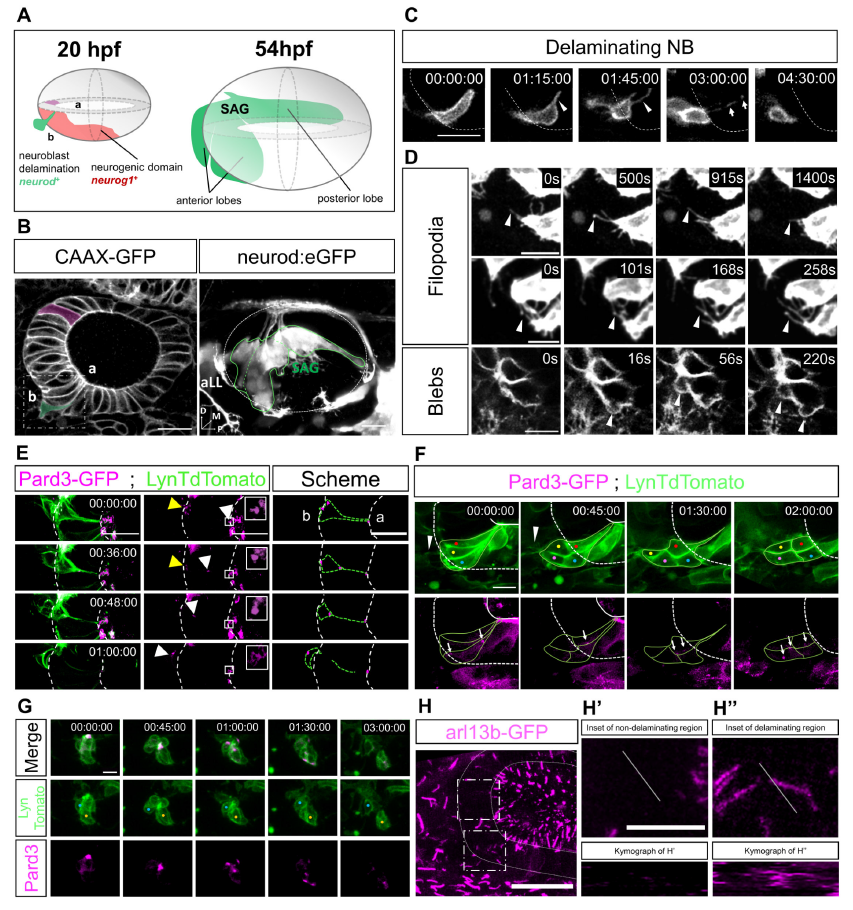
Single cell labeling and imaging of delaminating otic NB. A) Schematic representation of otic NB delamination in the neurogenic domain (red) at 20 hpf and SAG shape (green) at 54 hpf. B) Labeling of the otic epithelial membranes at 24hpf (left) and the SAG at 54hpf (right), the latter locating behind the otic vesicle (white dashed oval). Both images are lateral views. A NB (green pseudocolored) delaminating from the neurogenic domain displays a distinct shape compared to non-delaminating otic cells (purple pseudocolored). White square represents the region from where images in C and D panels are taken. Different SAG lobes are outlined with a green line. C) Single-labelled delaminating NB undergoing apical thinning (white arrowhead) while delaminating. NB exits neuroepithelia leaving membrane traces inside neuroepithelia (white arrows). D) NB from the neurogenic domain extend dynamic filopodia and blebs (white arrowheads) (n=7). Movie 2. E) Delaminating NB undergoing apical thinning and Pard3 relocation. Pard3-related change in polarity is observed by Pard3 movements from apical to basal domains concomitantly with membrane zippering (white arrowheads) and accumulation of Pard3 in basal domains (yellow arrowhead). Right panel represents a scheme of this phenomenon. F) Collective delamination of NB. NB extend filopodia in the collective front while delaminating (white arrowhead) and relocate Pard3 (white arrows) without losing its punctate pattern (n=7). G) After delamination, a group of contacting NB separate (blue and orange dots) concomitantly with Pard3 rearrangements (n=3). H) Otic vesicle non-neurogenic (upper dotted square) and neurogenic (lower dotted square) anterior domains, corresponding to insets H’ and H’’, respectively. Arl13b reporter is observed inside the neuroepithelia in the neurogenic domain, while absent in the non-neurogenic domain. A spatial section across time (white transverse line corresponding to kymographs) shows the presence of Arl13 reporter in the neurogenic region only (n=10). b: basal; a: apical; aLL: anterior lateral line ganglion; SAG: Statoacoustic ganglion. Anterior to the left, posterior to the right in all images. Scale bars 20µm in B and H, 10 µm in the rest.

To better characterize the cell shape changes during delamination in single NB, we fluorescently labelled the membranes in single and scattered NB by injecting lynTdTomato mRNA into one of the cells at 32-64 cell stage embryos. High spatiotemporal resolution in vivo imaging from 24 to 36hpf, the time window of highly active NB delamination, shows how a delaminating NB progressively acquires a triangular shape and concentrates the cytoplasm basally (Fig. 1C and E). During this cell shape rearrangement, the apical domain becomes narrower in a zipper-like process, to finally become a thin membranous filament (arrowheads in Fig. 1C). In the neural tube, it has been shown that the differentiating neurons detach from the apical membrane by a process of apical abscission, in which the cell is apically constricted and detached from the luminal membrane (Das & Storey, 2014). Individual labelling of otic NB allow us to also observe that when the basal cell body is mostly outside the epithelium, the attachment to the apical membrane is broken and the delaminating NB loses its apical contact, leaving some membrane debris behind (Fig. 1C white arrows, Fig. 1E, Movie 1). Using *in vivo* resonant scanning for higher temporal resolution (Fig. 1D) shows how predelaminating NB generate a high number of dynamic filopodia inside and outside the neuroepithelium as well as blebs at the basal cellular domain lining the basal lamina (Fig. 1D arrowheads from top to bottom rows, respectively. Movie 2). The basal lamina is disrupted at the neurogenic domain (unpublished results from the laboratory and Alvarez et al., 1989).

To ascertain whether apicobasal polarity is lost or rearranged during the delamination process. we assessed the localization of the apical determinants during delamination. For this, lynTdTomato mRNA was co-injected with the apical protein Pard3 mRNA at 32-64 cell stage. When analyzing Pard3 signal location in individual NB, it can be appreciated that a small fraction of Pard3 signal is kept at the abscised membrane that is left at the luminal area during the whole process (Fig. 1E, inset in central panel), but another fraction is detected regressing with the plasma membrane thinning edge (Fig. 1E, white arrowhead). In addition, accumulation of Pard3 is also transiently detected in the basal cytoplasmatic domain in early delamination (Fig. 1E, yellow arrow in central panel), suggesting a possible change in polarity (Movie1 and 3). When labeling is less mosaic, it can be observed several cells delaminating collectively (Fig. 1F). Some NB delaminate more dorsally (red, yellow and blue dots) and others more ventrally (magenta dot, absent in first frame). Delaminating NB produce dynamic filopodia (Fig. 1F, white arrowheads) and dynamically relocate puncta of Pard3 (Fig. 1F, white arrows).

Once NB are completely outside, they acquire a more mesenchymal and rounded shape. In a group of already delaminated NB contacting each other (Fig. 1G), a punctuated Pard3 pattern is initially observed at cell-cell contacts and is then redistributed as NB separate (Fig. 2C, blue and orange dots indicate cell bodies that separate through time). This suggests that in spite of a mesenchymal phenotype, polarity is reorganized and could be the driving force for NB dispersion (Fig. 1G and Movie 4). As observed in Fig. 1C and Fig.1E the delamination of NB spans for a period of 1,5-2 hours.

**Figure 2.**
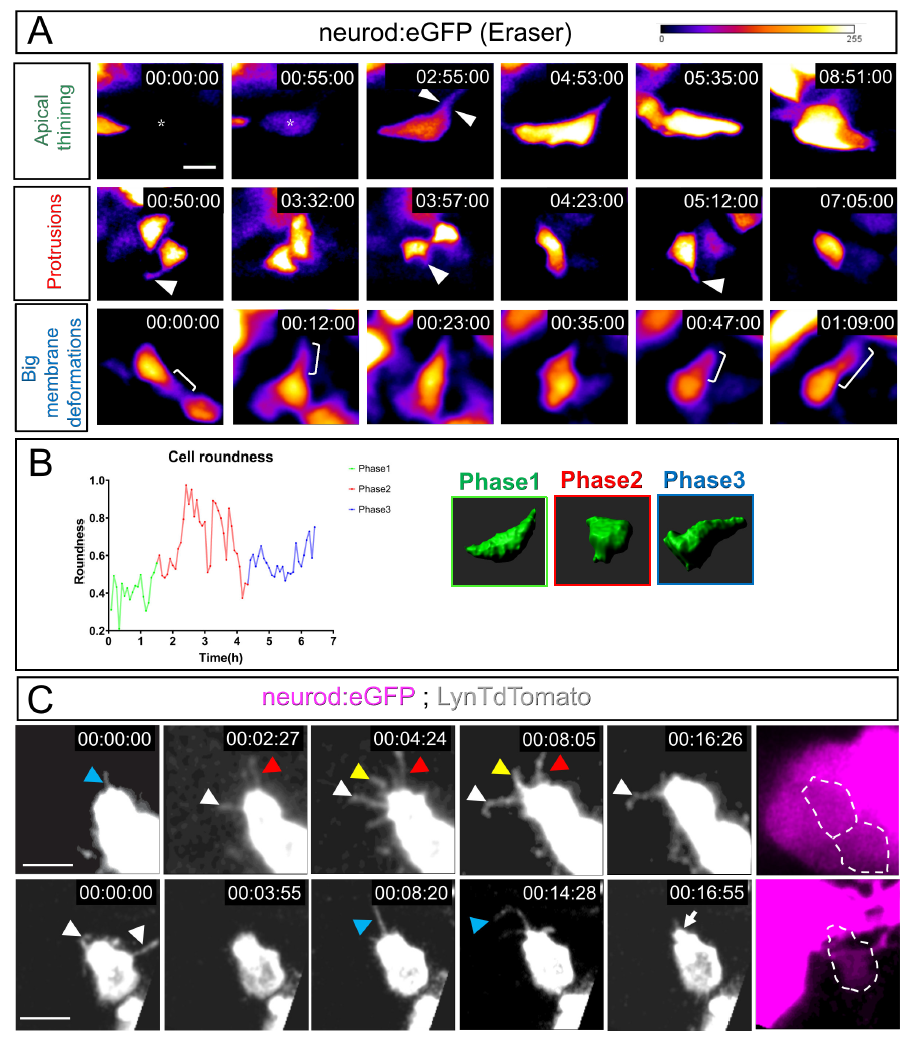
Delaminated otic NB dynamically change their shape, producing filopodia and membrane protrusions. A) Single cell labeling of delaminating NB with CRISPR Eraser in *Tg(neurod:eGFP)*. Delaminating NB undergo apical thinning and delamination (white arrowheads in top panel) right after start expressing *neurod* (asterisk). Followingly, they acquire a much more rounded, mesenchymal shape and produce membrane protrusions (white arrowhead middle panel). Finally, membrane protrusions precede big membrane deformations (white brackets). n=7. B) Example of roundness change of one NB through the different phases. Phases are color coded and a 3D IMARIS reconstruction is shown on the right. C) NB produce dynamic filopodia after delamination (colored arrowheads). Lower panel shows how a membrane protrusion becomes a much bigger membrane deformation (white arrow). Scale bars are 10 µm.

During the delamination of neural tube differentiating neurons, it has been shown that aPKC and the primary cilia are retained in the ventricle membrane when the cell suffers apical abscission (Das & Storey, 2014). In contrast, zebrafish retinal neuroblasts keep the primary cilia at the apical front of the retracting cell or either dismantle it only shortly before retraction during delamination (Lepanto et al., 2016). To study the localization of the primary cilia during otic NB delamination, we used the *Tg(actb2: arl13B-GFP)* line. In the non-neurogenic and neurogenic regions (upper and lower dotted squares respectively in Fig. 1H), it can already be appreciated how Arl13b staining is oriented towards the lumen in the non-neurogenic region but absent inside the epithelium, while present inside the neurogenic domain epithelium. This suggests that delaminating NB carry the primary cilia with them when delaminating, while non-delaminating cells keep Arl13b in the apical site (Inset Fig. 1H’ and Fig. 1H’’ and Movie 5). In a kymograph of the area (spatial section of these regions across time, white transverse lines in Fig. 1H’- H’’), Arl13b is found crossing the spatial section of delaminating NB but not crossing in non-delaminating region, again suggesting Arl13b locates inside the neuroepithelium only in the delaminating region (Fig.1H’ and H’’ kymographs).

Injection of mRNA at 32-64 cell stage results in few embryos with individual cells labelled. To increase the dataset of single NB labelled, we used what we have called CRISPR Eraser. In these experiments, a RNAguide against GFP is injected in embryos of the *Tg(neurod:eGFP)* reporter line that labels NB. Since Cas9 is highly efficient, most cells will have its GFP gene disrupted by Cas9 and only few NB retain cytoplasmic eGFP expression, either by escaping Cas9 targeting or because of efficient repair restoring eGFP genomic sequence (See Materials and Methods and Supplementary figure 1). In consequence, this creates an environment in which most of Neurod*^+^*cells of the developing SAG are eGFP negative and the remaining eGFP^+^ NB can be seen with high contrast.

In CRISPR Eraser embryos, NB cell bodies change from the elongated and triangular shape to a rounded cell body when exit the epithelium (Fig. 2A, white arrowheads in upper panel). Once delaminated, NB modify their roundness, extend filopodial protrusions that later on translate into larger membrane deformations (Fig. 2A, white arrowheads middle panel, Fig. 2C and Movie 6) and elongate (Fig. 2A, brackets in lower panel). During the whole span of the imaging of selected NB, the roundness has been quantified using FIJI and plotted in an example in Fig. 2B (Figure Supplementary2 shows more examples). At least three different phases of cell shape variation can be distinguished: an initial phase corresponding to apical thinning and delamination, a second phase of increased roundness and membrane protrusions and a final phase of elongation, presumably to engage into migration. IMARIS 3D reconstruction of a paradigmatic example depicts these three stages (Fig. 2B)

Individual labelling of NB and live imaging permits to highlight the remodeling events during delamination and outside the otic epithelium. Delaminating NB suffer a process of apical thinning and abscission but Pard3 and Arl13b components are not lost and relocate in the delaminated cell, suggesting that can help establish a new polarity front in NB. Moreover, NB extensively deform their membranes with large number of blebs, filopodia and larger protrusions, suggesting that might contribute to generate mechanical forces and/or establish cell communication events.

### Active migration of delaminated SAG neuroblasts is RhoGTPases-dependent

Massive delamination of zebrafish otic NB in the anterolateral region takes place between 17 and 30 hpf. Large groups of cells delaminate and position anterior to the otic epithelium (Hoijman et al., 2017). Thus, displacement of NB within the SAG can be driven mainly by new delaminating NB pushing on the previously delaminated cells. However, the dynamical changes in cellular shape and apicobasal polarity, together with the presence of filopodia observed in individual labelled NB, suggest that delaminated NB might display an active and directed migratory behavior.

Most of the cellular deformations produced by cells to actively migrate involve the activation of RhoGTPases. In particular, Rac1 RhoGTPase is implicated in directed migration and lamellipodia formation, Cdc42 in acquisition of migratory capacity and filopodia formation and Rho in stress fibers and rear contractility (Nobes & Hall, 1995; Reig et al., 2014; Shoval & Kalcheim, 2012; Yamao et al., 2015). To investigate the role of RhoGTPases in NB migration, we decided to manipulate this signaling pathway in NB. To this aim, we generated a CRISPR knock-in Gal4 line in the endogenous locus of the *neurod* gene which recapitulated the expression of the reporter transgenic line *Tg(neurod:eGFP)* (Supplementary figure 3). We then overexpressed different dominant negative (DN) and constantly active (CA) forms of RhoGTPases in the new *Tg(neurod:Gal4)* transgenic line.

**Figure 3.**
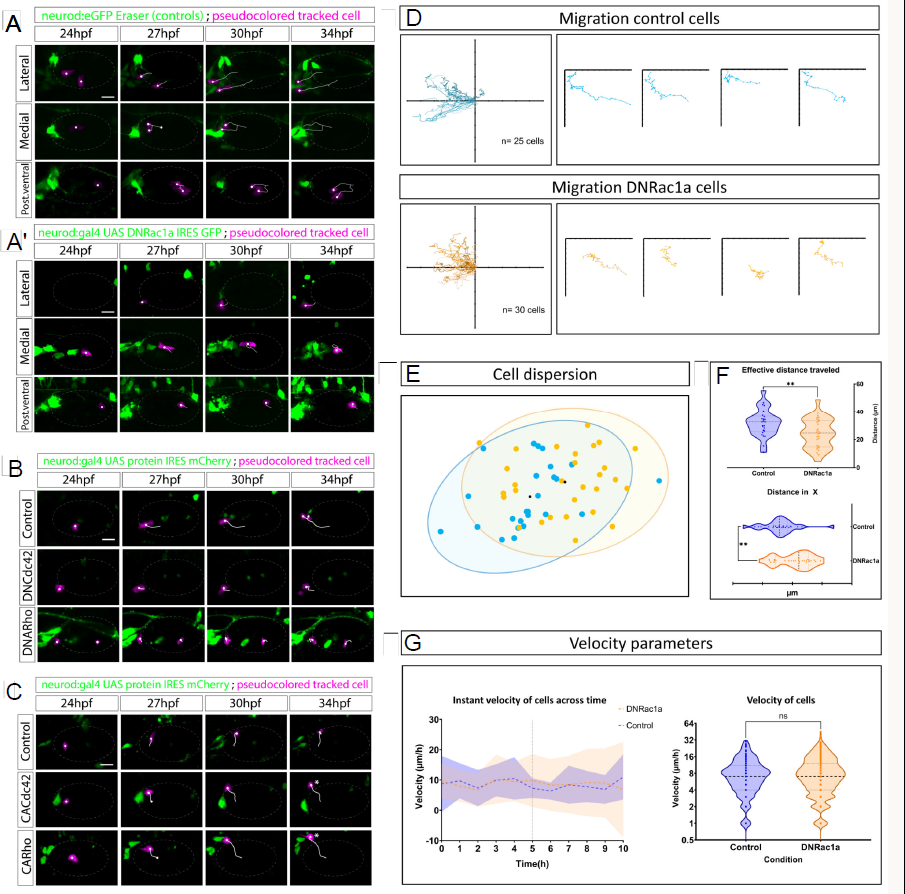
Otic NB engage into RhoGTPases-dependent active and directed migration. A and A’) Three NB migrating from lateral, medial and posteroventral regions in control wild-type conditions under CRISPR Eraser upon *Tg(neurod:eGFP)* or overexpressing DNRac1a (magenta pseudocolored cells, tracks in white). Otic vesicle is depicted with a dim white dashed line. Scale bars are 20µm. Associated Movie 7. B) Migratory pattern under DNCdc42 and DNRho. DNCdc42 and DNRho recapitulate the effect of reduced directed migration. C) In contrast, CACdc42 and CARho show similar or even enhanced migration compared to control cells (*). D) Summary of tracked control and DNRac1a-expressing cells using DiPER.Tracks are normalized at the origin according to (Breau et al., 2017; Gorelik & Gautreau, 2014). Control cells migrate more distance and more directionally than DNRac1a cells. E) 95% confidence interval for dispersion of cells at endpoint of the timelapse after normalization of tracks at origin, in control (blue) and DNRac1a (orange) conditions. F) Effective distance migrated is significantly (**) compromised in DNRac1a compared to controls, measured as the length of a straight line between start and end point of a given cell irrespectively of their particular migratory path in the recordings. Distance covered in the X axis is also compromised (**). G) Instant velocities (space covered/time between frames) of delaminated NB across time. Migratory capacity of cells seems little affected since the mean (blue and orange dashed lines) of both control NB and DNRac1a NB is the same, although dispersion of standard deviation increases towards the end of the recording in DNRac1a condition (orange area) versus controls (blue area). Black vertical dashed line indicates half of the recording time. For detailed tracked velocities see Supplementary figure 4. Otic vesicle is depicted with a white dashed line. For DNRac1a analysis, phenomenon observed in n= 25 control cells and n= 30 DNRac1 cells, from 7 and 14 embryos, respectively. Scale bars are 20µm. See Supplementary figure 4 for individual cell migratory profile. With respect to the Cdc42 and Rho1 experiments, control cells of the DN condition migrate normal in 2/2 cases; DNCdc42 cells migrate normal in 4/7; DNRho cells migrate normal in 1/4 cases. Control cells of the CA condition migrate normal in 5/5 cases; CACdc42 cells migrate normal in 3/6; CARho cells migrate normal in 1/4 cases. Controls are injected with UAS: mCherry plasmid. All embryos are siblings from the same batch of injection. Otic vesicle depicted with dim grey dashed ellipse. Tracked cell is pseudocolored in magenta and tracks in white. Scale bars are 20 µm. Ticks in X axis represent increase every 10µm Associated Movie 8.

We first overexpressed the dominant negative (DN) form of Rac1 by injecting at 1-cell stage embryos the UAS: DNRac1a-F2A-GFP Tol2 construct (Hanovice et al., 2016). We then tracked, over a period of 10 hours, control cells under CRISPR Eraser condition and the Rac1 inhibited cells expressing GFP (Fig. 3A versus Fig4. 3A’ and Movie 7). Compared with control cells, DNRac1a cells showed a reduced migration distance irrespectively of their initial position (lateral, medial or posteroventral; compare white tracks from pseudocolored magenta cells in Fig. 3A versus 3A’ and D,). All tracks are plotted in Fig. 3D using DiPER (Gorelik & Gautreau, 2014), as well as 4 individual ones as examples (n= 25 control cells and n= 30 DNRac1 cells, from 7 and 14 embryos, respectively). Fig. 3E represents a 95% confidence interval of 2D spatial dispersion of NB considering the final position of every NB in control (blue dots) and DNRac1a (orange dots) conditions after normalizing the tracks to origin (Breau et al., 2017; Gorelik & Gautreau, 2014). In summary, Fig. 3E shows the covered area (ellipses) or dispersion of NB in control vs DNRac1a conditions after assuming they all depart from the same point (See 2D Dispersion analysis in Materials and Methods section). Non-overlapping regions and different ellipse centroids (black dots Fig. 3E) reveal differences in the dispersion of NB between conditions. Statistically significant representations of such differences are the reduced distance in X axis and the reduced effective distance traveled of DNRac1a NB versus control NB (Fig. 3F, ticks in X represent 10µm increase). The effective distance traveled by cells is the straight line from initial to final position.

Comparing quantified velocities across time (Fig. 3G left graph and Supplementary figure 4), DNRac1a-expressing NB are similar to control cells. However, migrated distance is reduced in DNRac1a conditions (Fig. 3E-F). This indicates that the directionality of migration in DN-expressing NB is impaired. In summary, DNRac1a NB still move and at similar rate velocities to controls but lose the persistence in directed migration. We extended the analysis by abrogating (DN) and increasing (CA) the function of the RhoGTPases Cdc42 and Rho1 in otic NB (Fig. 3B-C, Movie 8). In these conditions, we found that NB have a reduced or recovered migratory capacities, respectively (Fig. 3C, asterisk).

**Figure 4.**
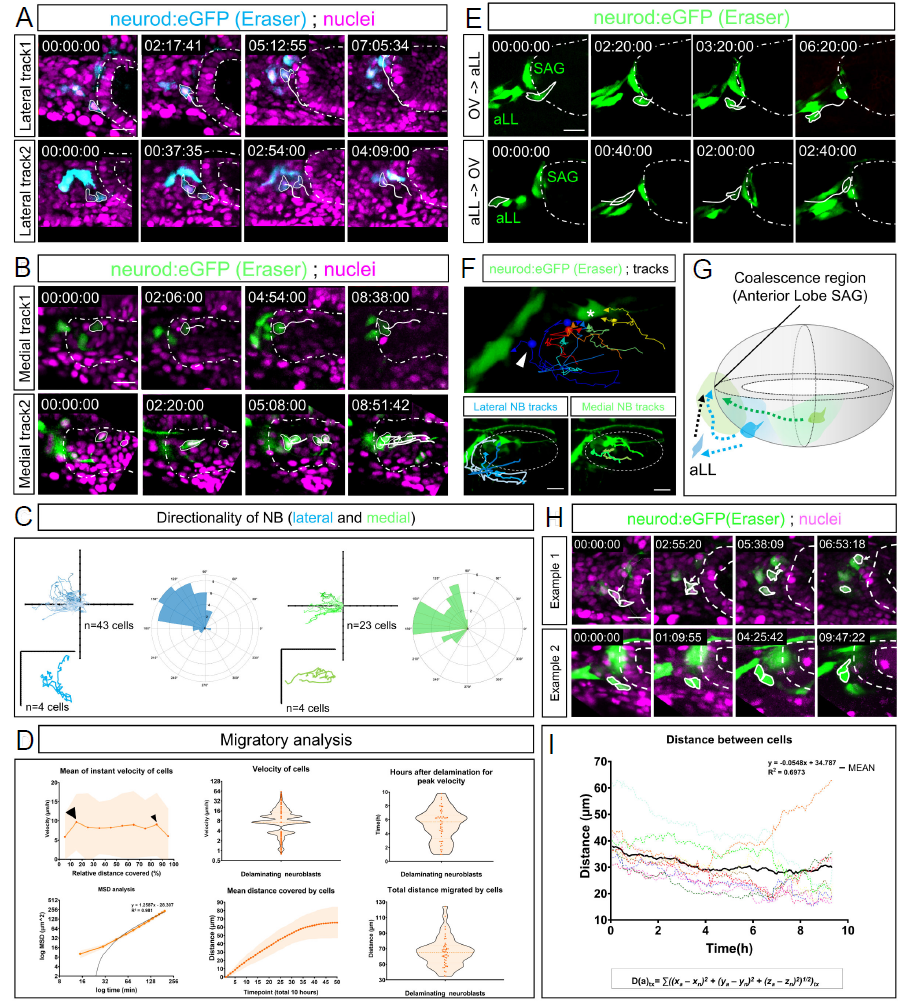
Delaminating NB directionally migrate towards a common coalescence region. A) Two tracks (white lines) of delaminated NB from lateral domains of the otic neurogenic region. B) Two tracks (white lines) of delaminated NB from medioventral domains of the otic neurogenic region. Otic vesicle contour depicted inside white dashed line. Scale bar is 20µm. See Supplementary figure 5 and associated Movie 9 and Movie 10. C) Summary tracks using DiPER normalized at origin and directionality rose plots (cyan for lateral and green for medioventral) of delaminating NB (Breau et al., 2017; Gorelik & Gautreau, 2014) showing migration of NB is directed. Ticks in axes represent 10µm increase. D) Graphs of quantified data of NB migratory properties. For a separate analysis of lateral and medial delaminating cells Supplementary figure 5. E) Interchange between aLL and SAG and vice versa. F) Tracking several NB in the same embryo shows aggregation towards a common region (*) irrespectively of NB origin inside the otic neuroepithelia. White arrowhead indicates an otic NB attracted to aLL. G) Graphic summary. H) Qualitative examples of non-collective and collective migration. I) Plotting several cases summing the distance of one NB with respect of its neighbors across times shows a negative regression, which is indicative of distance shortening between cells and thus, aggregation. Otic vesicles depicted inside white dashed lines. Anterior always to the left and posterior to the right. Scale bar 20 µm.

Altogether, our data blocking and activating RhoGTPases reveal for the first time that RhoGTPases pathway is required for a directed and active migration of delaminated otic NB. This suggests that otic NB engage into active and complex behaviors to organize the SAG, rather than undergoing passive and bulk organization.

### NB display a characteristic migratory profile to a coalescence region

Under the findings that otic NB actively migrate, we then wondered about which were the migratory paths followed by NB. Using CRISPR Eraser in *Tg(neurod:eGFP)* and photoconverted cells in *Tg(neurod:kikume)* background, we tracked delaminating NB starting at their initial point of delamination in the neurogenic domain for a period of 10- 12 hours from the delamination peak of 20-24hpf until 34-36hpf. NB delaminating from the anterolateral neurogenic domain follow a longer migratory path and move from lateral to medial (Fig. 4A, cyan color coded; Movie 9). On the other hand, NB delaminating from posteromedial regions migrate anteriorly to the same region between the OV and the HB (Fig. 4B, green color coded; Movie 10). Wind rose plots of the directionality of paths confirm this directed anteromedial migration of delaminated NB (Fig.4C, color coding preserved). The tracking of many migrating NB (n = 66 [43 lateral + 23 medial] cells from 40 embryos) highlights their migratory directionality towards a region just anterior and medial to the otic vesicle and attached to the hindbrain wall at rhombomere 4 (Fig. 4F, asterisk; Fig. 4G). Migration and coalescence to this region contributes to the growth of the anterior lobe of the SAG (Fig. 4G). We named this region the coalescence region. Although it has been suggested that medial delaminated NB position in the posterior lobe (Dyballa et al., 2017), tracking of medial delaminated NB show that also migrate and coalesce anteriorly.

Quantification of the time elapsed from delamination until cell reaching the coalescence region, which normally coincides after a rapid movement of peak velocity, indicates that it takes around 6h (Fig4.D, smaller black arrowhead and Fig. 4D, upper right corner panel; Supplementary figure 4C, 4C’ and 4E black arrowheads).

When the mean of instant velocities of NB against normalized distance covered is plotted, a slight increase in velocity just right after delamination is appreciated to then display a fluctuating walking behavior (Fig. 4D, thick black arrowhead). NB interchange steady moments with pulses of movement, which is indicative for the previously reported active migration of NB and coinciding with the three different phases of behaviors described in Fig 2A-B. Finally, they increase the velocity until they reach a region where they stop and coalesce (Fig. 4D, thin black arrowhead; Supplementary Figure 5 distinguishing lateral and medial delaminating NB behaviors). Quantified maximum velocity of migration is 61’2µm/h (Fig. 4D), maximum distance covered is 124µm and mean distance migrated 66µm. MSD analysis with DiPER (Gorelik & Gautreau, 2014) of our data shows a slope (black line) of the linear fit that is greater than 1 (α > 1), which is indicative of directed motion. For further detailed and separated analysis between more lateral and more medial NB see Supplementary figure 5.

**Figure 5.**
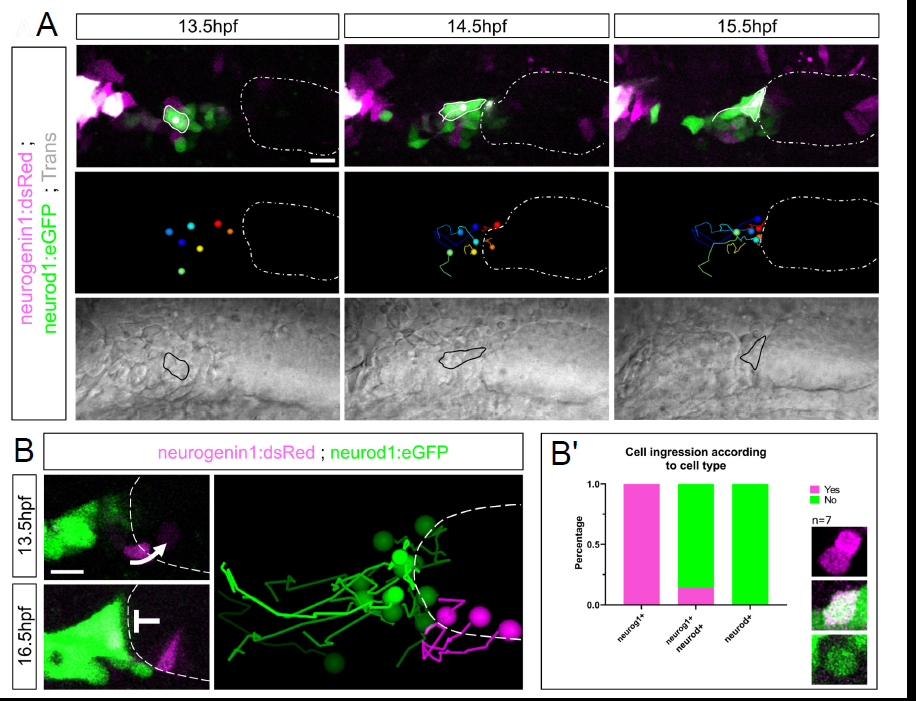
Pioneer SAG neurons of extraotic origin initially populate the coalescence region. A) Extraotic pioneer SAG neurons are specified anterior to the otic placode (white dashed oval) and posterior to the trigeminal ganglion (magenta signal in left corner) at around 13hpf. Pioneer SAG neurons migrate and initially populate the coalescence region in anterior locations of the otic placode. B) Pioneer SAG neurons are neurod+ cells that do not ingress into the otic epithelium. B’) Quantification of number of cells ingressing or not depending on gene expression. Scale bars in A is 20µm, 10µm in B.

Surprisingly, in some instances, we also observed otic delaminating NB being able to incorporate into the anterior lateral line (aLL) ganglion and vice versa, suggesting that both neuronal tissues are plastic receiving and sending neuronal progenitors (Fig. 4E; Fig. 4F white arrowhead) whose fates are not fully determined yet.

We previously observed that delamination of NB can be collective. We then wondered if migration of NB was also collective. Collective migration is defined by a group of cells that keep cell-cell contacts, have group polarization and exhibit a coordinated behavior relevant for the proper organization of the tissue (Le et al., 2022). To address this question, we observed in several cases two touching NB and their neighbors. As shown in Fig. 4H, delaminated NB can either migrate non-collectively (Fig. 4H, example 1 white arrow; Movie 11) interrupting their physical contact and NB separating each other. In other cases, two touching cells migrate together (Fig. 4H, example 2, Movie 12), maintaining their physical contacts during the whole imaging period. Therefore, the data suggests that NB migration within the SAG does not require permanent NB cell-cell contact per se.

Nevertheless, our previous results indicated a common pattern of migration to the coalescence region performing a migration in streams. When analyzing and plotting the distance between several NB against the rest of the neighbor cells across time, a negative linear regression appears, confirming NB coalescence as a common feature for migrating NB (Fig. 4I, black line).

In summary, NB delaminating from different regions of the neurogenic domain migrate and aggregate around a particular region, which we called the coalescence region. The coalescence of NB in this region gives rise to the anterior lobe of the SAG (Movie 13).

### Pioneer SAG neurons populating the coalescence region are required for NB migration and SAG organization

The coalescence region where delaminating NB aggregate is populated by bright *neurod^+^* expressing cells. To assess the origin of these cells, we performed time-lapse analysis at 13.5hpf, before otic delamination begins. A group of few scattered cells expressing either *neurogenin1*, *neurod* or both are detected anterior to otic placode territories and posterior to the trigeminal ganglion (Fig. 5A). These cells migrate to anterior regions of the otic vesicle (Fig. 5A white dashed oval) and apposed the 4^th^ rhombomere wall by 16hpf, populating the location that will later become the coalescence region. From this group of cells, the subset of *neurogenin1^+^-*only cells, observed with the reporter line *Tg(neurog1:dsRed),* ingress into the otic epithelium as shown previously in (Hoijman et al., 2017), whereas the subset of *neurod^+^* cells remain outside (Fig. 5B-B’). This latter group was called pioneer SAG neurons, since they are the first ones populating the prospective SAG ganglion.

To address whether pioneer SAG neurons have a role in migration and coalescence of NB, at 16hpf we first photoconverted these cells in our *Tg(neurod:kikume)* line from green to magenta and then ablated them with 2-photon microscopy (Fig. 6A). Attending to different experimental conditions, whether pioneer SAG neurons remain unablated, partial (Fig. 6B, asterisk) or total ablated, various SAG defects are found at later stages (Fig. 6B, blue dashed line indicates SAG shape in Z projections of images). In ablated embryos, SAG shape is aberrant and the number of cells populating the SAG is reduced at 24hpf (Fig. 6B and 6C’’’, n = 23 control cells, n = 36 ablated condition cells, from 4 embryos and 10 embryos, respectively). At 34hpf, the formation of the posterior lobe is abrogated (Fig. 6B, white arrowheads). SAG shape is altered because lack of pioneer SAG neurons causes altered directionality patterns of migration and altered dispersion of NB, found after tracking again, using DiPER, individual NB when SAG pioneer neurons are missing (Fig. 6C’-C’’, Movie 14; Supplementary figures 6-8). Fig. 6C’’ shows, as in Fig. 3E, a 95% confidence interval of dispersion of NB at final position when normalized at origin between control (green) and ablated (magenta) conditions. Non-overlapping regions evidence differences in NB dispersion, which are statistically significant according to Y axis distance (Fig. 6C’’). Although NB migrate less directionally, NB in ablated condition are capable of migrating faster (Fig. 6C’’’ upper panels). Moreover, absence of pioneer SAG neurons affects the number of NB that form the SAG and consequently, SAG volume is reduced (Fig. 6C’’’ lower panels and Fig. 7).

**Figure 6.**
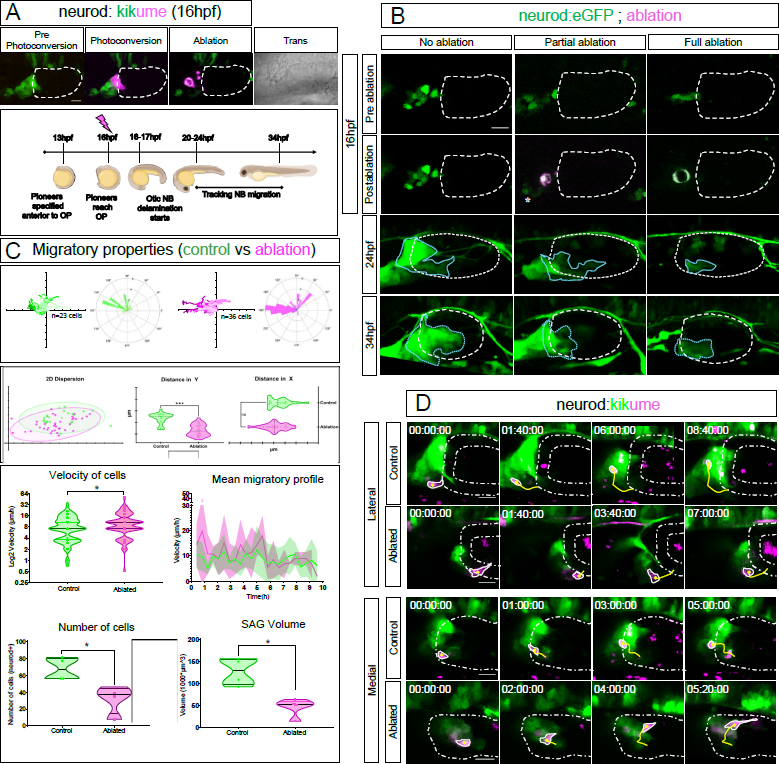
Pioneer SAG neurons have a role on organizing the SAG. A) Scheme of experimental design. At 16hpf pioneer SAG neurons are photoconverted and subsequently photoablated. The absence of magenta photoconverted cells at later stages indicates complete cell ablation. SAG development is imaged from 20- 24hpf to 34hpf. B) After partial (*) or total ablation of neurod+ cells anterior to the otic placode at 16hpf it can be observed already at 24hpf an altered shape of the SAG and an apparently reduced number of NB. At 34hpf, the formation of a posterior lobe is abrogated in the ablated conditions (white arrowhead and Supplementary figure 7) (experiment replicated 3 times. n number of embryos; n control = 4+3+6; n ablated condition = 4+3+11). C) Migratory properties of otic NB under control (green) or pioneer SAG neurons ablated (magenta) conditions measured with DiPER (Breau et al., 2017; Gorelik & Gautreau, 2014). C’) NB migration directionality and pathway is compromised in pioneer SAG neuron ablation condition compared to controls. n control cells = 23, n ablated condition cells = 36, from 4 and 10 embryos, respectively to each condition. Every tick in axes represent 15µm increase. C’’) 95% confidence interval of NB dispersion according to last timepoint position when tracks are normalized at origin in control (green dots) versus ablated (magenta dots) conditions. Non-overlapping regions evidence the variance in dispersion, which is significant in the Y axis (***) while non-significant in the X axis (ns). Ticks in axes represent increase every 10µm. C’’’) NB are able to migrate faster (*) in the ablation condition. Mean migratory profile is not majorly affected, except that dispersion of velocity is increased in the ablated condition at initial stages of migration (Supplementary figure 8 shows detailed individual migratory profiles). Number of cells populating the SAG is significantly reduced (*) in the ablation condition, more than the number of cells ablated (Supplementary figure 6A) and consequently SAG volume (Supplementary figure 7). N number of embryos for this section (C′′′) is 4 both for controls and ablated. D) Two examples of migration of NB (photoconverted magenta cell, yellow tracks) between lateral and medial regions of the neurogenic domain. Otic vesicle depicted between white dashed line. Total control embryos n = 6; total ablated embryos n = 11 for this section (D). Scale bars are 20 µm.

**Figure 7.**
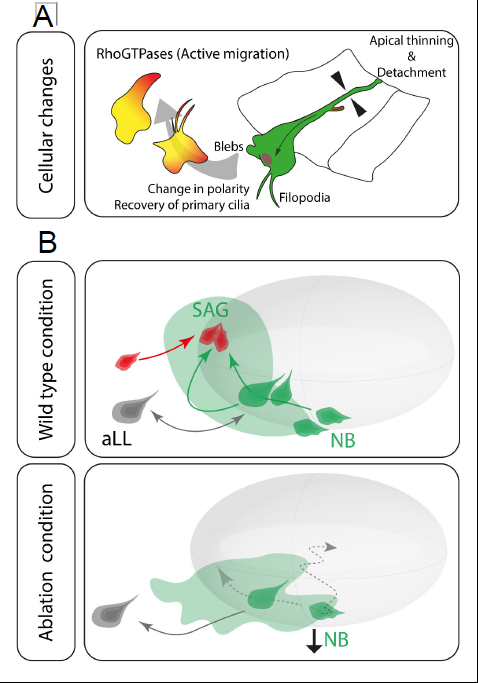
A) Summary of cellular behaviors of otic NB addressed in this study. B) Schematic drawing summarizing effects derived from the lack of pioneer SAG neurons (red).

Altogether, our results point towards a fundamental role of pioneer SAG neurons in recruiting delaminated NB and thus orchestrating the shape and growth of the SAG (Fig. 7).

## Discussion

Morphogenetic events of cranial sensory ganglia development at the cellular level are little understood in comparison to other model systems such as epithelial tissues. In this work we provide, for the first time with cellular resolution and in vivo analysis, the cell behaviours underlying NB delamination, migration and coalescence of the statoacoustic ganglion.

### NB otic delamination as an EMT process

Whether otic NB delamination follows an EMT program has been under debate. Some authors have suggested that sensory neuron delamination in cranial placodes is not an EMT event (Graham et al., 2007; Schlosser, 2008, 2010). It was reported that Snail family of core-EMT transcription factors and RhoGTPases (specially RhoB) are not used in this process (Graham et al., 2007). Moreover, Sox9/10, important EMT regulators, were found to be absent in most *Xenopus* placodes (Schlosser, 2008, 2010). Additionally, migratory capacity of cranial placode NB seems diminished compared to neural crest (Schlosser, 2008). However, other evidences suggest the contrary, where published gene expression data and our imaging data points towards a delamination process in the inner ear with EMT characteristics.

Otic delaminating NB express some EMT-associated genes such as *sox9a/b* (Yan et al., 2005), *sox10* (Chiang et al., 2001; Dutton et al., 2009; Piloto & Schilling, 2010), the Spemann organizer gene *goosecoid* (Kantarci et al., 2016), *snail1b* (Léger & Brand, 2002; Zecca et al., 2015) and *cdh6* (LF, unpublished results and (Hans et al., 2013)), which is known to be important for NCC migration (Clay & Halloran, 2014). Together with gene expression changes, EMT is accompanied by large cell shape rearrangements, acquisition of extensive migratory capacity, apicobasal polarity changes and cell adhesion disassembly (Yang et al., 2020). Finally, otic NB are still not postmitotic (Camarero et al., 2003; Hoijman et al., 2017), meaning delamination is not strictly of neurons, as it is in other placodes (Graham et al., 2007).

We show here the changes in cytoarchitectural dynamics, apical determinant and primary cilia relocalization during otic NB delamination. The sequence of cellular cell shape changes involves an apical thinning, with most cytoplasm concentrating basally, an eventual apical detachment from the lumen with some membrane remaining’s, resembling what was described in neuronal neural tube delamination (Das & Storey, 2014); extensive basal blebbing, lateral and basal filopodia extensions and final NB translocation out of the epithelium. Blebbing has been shown to be an important mechanism to facilitate basal lamina removal and EMT in zebrafish NCC (Berndt et al., 2008; Font-Noguera et al., 2021; Paluch & Raz, 2013). However, it is not clear yet in our tissue whether blebbing occurs as an active property for facilitating delamination/migration or a passive event due to forces from neighbouring cells and loss of basal lamina and membrane integrity (Schick & Raz, 2022).

When imaging apical protein Pard3 localization during otic delamination, Pard3 is partially kept in the apical membrane remaining’s after abscission, while a different fraction travels basally in the hinge point of apical cytoplasmatic thinning, suggesting it might provide information for the apical thinning process. In our system, compared with the laterality organ of zebrafish (Pulgar et al., 2021), partial delamination does not occur, but once NB are out of the epithelium, Pard3 is dynamically rearranged while retained, similarly as in in the zebrafish heart trabeculation (Jiménez-Amilburu et al., 2016). Pard3 maintenance in pseudo-mesenchymal NB suggest that Pard3 could establish a new polarity axis in migratory cells or inform where neurites must grow, as shown in some mechanosensory neurons (Lee et al., 2021). In NCC, relocalization of the centrosome on the other side of the nucleus during EMT results in a polarity reversal and is responsible for cell dispersal when become mesenchymal (Burute et al., 2017). Moreover, Pard3 localization in cell-cell contacts during NCC migration promotes microtubule disassembly (Moore et al., 2013).

In chick, an actin-based apical constriction leaves an apical abscised part containing the primary cilia membrane but centrosome remains in the delaminating cytosol (Das & Storey, 2014; Kasioulis et al., 2017). In our delaminating NB we observe internalization of the primary cilium. The fact that the primary cilium is not dismantled nor kept on the abscised membrane might favour rapid reestablishment of polarity once delaminated NB are outside. Whether there is a passive apical membrane stretching and shearing while recovering apical proteins and primary cilium or there is an active apical cell-autonomous cytoskeletal rearrangement was not studied here. It is known than Cdh6 and RhoA in apical domains lead to the apical constriction in NCC engaged into delamination *in vivo* (Clay & Halloran, 2014).

In summary, data from our experiments and literature suggest that otic NB undergo an EMT process when delaminating.

### NB as active entities to organize the SAG

Once outside the otic neuroepithelia, otic NB acquire a much variable and rounded shapes. We have shown delaminating NB actively engage into complex migratory behaviours in a RhoGTPases-dependent manner. This suggests that NB migratory capacity is more constrained by neighbouring tissues and signalling than by low migratory capacity (Graham et al., 2007; Schlosser, 2008). Moreover, it has been shown that NCC touching the SAG maintain SAG aggregation but are not implicated in giving migratory cues to NB (Sandell et al., 2014; Zecca et al., 2015). In NCC, reducing Rac1 and consequently reducing lamellipodia extensions affects NCC migration while delamination still occurs. This indicates that these protrusive extensions are required to generate a migratory force in NCC, and thus large protrusions might be required in our tissue for the same purpose.

We have also seen NB can migrate either collectively or non-collectively. In a collective migration, cells can move cryptically within the cluster (Reig et al., 2014) but the overall migration is robust. In NCC tissue, E-cadherin helps maintaining stream migration while Pard3-N-Cadherin interactions avoids crowding by contact-inhibition of locomotion (Clay & Halloran, 2014; Moore et al., 2013; Priya & Yap, 2015; Scarpa et al., 2015). If a temporarily regulated and partial switch in adhesion molecules is necessary to maintain some sort of collective migration for keeping tissue cohesiveness remains to be studied. We do know that itg5a is required for correct aggregation of epibranchial placodes (Bhat & Riley, 2011), maintaining tissue segregation from other placodes and cohesiveness within the tissue.

Another key finding of this work is that delaminated NB engage into active migration towards a coalescence region, irrespectively of their origin location inside the neurogenic domain and their delamination position. Thus, when NB delaminate they do not remain in the same location, but rather migrate and coalesce anteriorly. This suggests that the SAG organizes in a much more complex manner than assumed according to genetically position information.

### The Posterior Placodal Area providing Pioneer neurons as organizing centres for SAG assembly

Neurogenic cells from the Posterior Placodal Area (PPA) were known to exist but thought to contribute exclusively to aLL or the otic placode (Andermann et al., 2002). Here we show for the first time how PPA neuronal specified cells can contribute also to the SAG. These cells have a pioneer role over otic delaminated NB and could also have a pioneer role in the aLL. In addition, we also observe that, in some cases, delaminated NB can integrate into the aLL and viceversa. Thus, at very early stages, neurogenic cells of the PPA are still not fully committed and can be interchangeable between aLL and SAG, in accordance to the hypothesis put forward in Abelló & Alsina, 2007. The signals that help pioneer SAG neurons to position anterior to the otic vesicle are unknown. The SDF1/cxcr4 system has a role on the lateral line primordium migration, however SDF1 has not been reported to be expressed in the otic placode. Molecules from the hindbrain could also play a role driving pioneer SAG neuronal populations migration. Finally, chase and run from NCC-placodal neuron interactions (Theveneau et al., 2013) is discarded due to time window constrains as pioneer SAG neurons migrate before NCC invade the tissue. NCC would have a later role in maintaining SAG cohesiveness as suggested (Begbie & Graham, 2001; Steventon et al., 2014; Zecca et al., 2015).

Pioneer neurons and tracts were discovered in seminal studies in *Drosophila* (Harrison, 1959) and recent studies show to be required for other neuronal cell body correct positioning (Karkali et al., 2023). In zebrafish, a group of pioneer neurons is required for correct migration of later facial branchiomotor neurons (Wanner & Prince, 2013). Here, we provide novel evidence on pioneer SAG neurons of non-otic origin that by positioning adjacent to the otic vesicle have also a role on the migration of delaminated otic NB and the organization of the anterior SAG lobe.

Our current work identifies a group of pioneer neurons with a role in otic NB migration and coalescence depending on pioneer SAG neurons. This knowledge will expand our understanding on the development of cranial ganglia, in which pioneer neurons prefigure the definitive architecture and/or location of a neuronal tissue.

## Materials and Methods

### Fish maintenance and husbandry of transgenic lines

Zebrafish embryos and adults were maintained and handled according to standard procedures at the aquatic facility of the Parc de Recerca Biomèdica de Barcelona (PRBB), in compliance with the guidelines of the European Community Directive and the Spanish legislation for the experimental use of and as previously described (M., 2000). Stable transgenic lines were kept by means of alternate outcross with WT (AB/Tü) and incross, generation after generation. Expansion of the lines were done every 1,5-2 years. Embryos were kept under dark conditions at a temperature of either 23 or 28’5°C in Danieau’s solution.

For this study we used the following lines:

*TgBAC(neurod:eGFP)^nl1^* labeling specified neuroblasts and neurons (Obholzer et al., 2008); *<u>Tg(neurod:Gal4</u>)* line generated in the lab in which Gal4 expression is driven by the neurod1 promoter, insertion by CRISPR Knock-in following the protocols from (Auer, Duroure, Cian, et al., 2014a; Auer, Duroure, Concordet, et al., 2014; Auer & Del Bene, 2014; Kimura et al., 2014) using the gbait plasmid from (Kimura et al., 2014); *<u>Tg(neurod:Gal4; UAS: H2A-GFP)</u>* combining the Tg(neurod:Gal4) with injection of the Tol2 plasmid [UAS-H2A-GFP], kindly provided by Dr.Jeroen Bakkers’ Lab (Strate et al., 2015); *<u>Tg(neurod1:Kikume)</u>* generated in the lab by injecting a plasmid containing 5kb neurod promoter upstream the photoconvertible protein kikume. Plasmid kindly provided by Dr. Katie Kindt’s Lab; *Tg(neurogenin1:dsRedE ^nl6^)* labeling early specified neural progenitors (Drerup & Nechiporuk, 2013); *<u>Tg(actb2:H2A-mCherry)</u>* as a pan-nuclear marker, line provided by Dr. Esteban Hoijman at Dr. Verena Ruprecht’s Lab; and *Tg(ubb:arl13b-EGFP)* labeling the primary cilia related protein Arl13b (Austin-Tse et al., 2013).

Embryos used later than 34 hours post-fertilization were kept transparent by soaking them in embryo medium (Danieau’s Solution) with 1% 1-phenyl-2-thiourea (PTU) (Sigma) to inhibit pigment formation (M., 2000). This treatment did not affect development in controls. Embryos were staged as previously described (Kimmel et al., 1995). In all experimental conditions the embryos of control and experimental conditions were siblings except when comparing DNRac1a with CRISPR Erasers.

### Microinjection

Long and very thin injecting needles were done in an electrophysiology puller (Sutter instruments model P-97) with the following protocol: P=200; HEAT=566; PULL=90; VEL=70; TIME=80. The tip of the needle was bevel broken by using forceps. Needle was loaded with injecting solution. Embryos were injected with 1 or 2nL into the cell or yolk at 1 cell stage embryos. Injections of mRNA ranged from 50 to 250ng/µL. RNA synthesis as described according to guidelines from commercial house (#AM1340 Invitrogen). sgRNA for GFPbait as described in (Gagnon et al., 2014).

Mosaic injections at 32-64 cell stage were performed in the central cells according to fate map analysis (Strehlow et al., 1994; Strehlow & Gilbert, 1993) in which the probability of getting our tissue of interest labeled is maximized.

### DAPI Staining and cryosections

From a stock at 5mg/mL we performed a 1:500 or 1:10.000 dilution in PBS with 0’1%Tween-20 (PBT 0’1%) for whole zebrafish up to 24hpf or slides, respectively. Embryos or slides were submerged in the solution for 5 minutes and washed for 5 minutes in PBT 0’1%. In slides, specialized slides were used (StarFrost Objektträger Knittel glass). Cryostat sections 20µm thick. Protocol of cryosection as described in (Taberner et al., 2020).

### FIJI processing

For visualization purposes only, images were non-linearly processed with FIJI plugins subtract background (50 pixels radius), smooth and/or gaussian 3D blur with X, Y, Z values of 1 or 2. Images were also linearly modified with brightness/contrast tool and median filter with the same values within the same experimental batch. 3D drift correction was made according to nuclei, SAG and/or transiluminated otic vesicle with the “3D drift correction” FIJI plugin (Movie 15). Activation of enhanced drift correction options. Images are either Z projections from a stack or single planes.

### Confocal live-cell imaging

Anesthetized zebrafish (42µL tricaine per mL of water, from a stock of 400mg tricaine powder in 100mL H2O) were mounted on on a 35 mm Ibidi μ-dish (#81156 Ibidi) in 1% low melting point agarose 45° tilted from completely dorsal. Imaging of embryos was performed on an inverted Leica SP8 confocal system equipped with a 488 nm Argon-ion laser (LASOS, 488 nm laser power 2.5 mW at back focal plane at 100% laser power), 405, 561, 633 laser diodes, motorized xy and z-galvo stage, 1 HyD and 2 PMT detectors and a HC PL APO 20x/0,75 IMM and CS2 objective (#506191 and #506423). The 488nm laser excitation depended on the experiment but was always kept across samples for the same and related experiments. For in vivo experiments it ranged from 0,1 to 10%. For fixed embryos, it could increase up to 80%. Gain normally ranging between 600-900Hz and off-set was never higher than 0,1%. Fluroescence excitation at 488nm was collected between 500 and 543nm; and fluorescence excitation at 561nm was collected between 589 and 621nm. Using Leica LUT setting2 sets a high-low mode, which is used for setting the pixel saturation limits (background pixel value 0 = green, saturated pixel value 255 = blue).

Frame 1024×512 or 1024×1024 at 8-bit. Scan speed ranged from 400 to 1000Hz (normally 600Hz). Z step sizes are system optimized according to Leica settings (normally a HC PL APO 20x/0,75 IMM #506191 give a z-step size optimized of 1.19µm with pinhole in airy1). Software zoom factor was set at 2,5x for most of experiments. Bidirectional scanning was activated. Line average and frame average either 1 or 2 was used. Multi-position (Mark & Find) experiments included up to 35 embryos separated by a time frame no longer than 20 minutes, except where otherwise mentioned. To avoid cross-talk between fluorophores sequential line acquisition was used. Cross-talk was in silico checked using FP base (https://www.fpbase.org/spectra/).

Tissue drift was considered and corrected. Otic tissue moved anteriorly at a rate of or 4- 10µm/h. Videos of up to 12 hours considered a total anterior drift of 120µm maximum.

### Resonant scanning

To maximize temporal resolution time-lapse imaging of filopodia, the resonant mode (8000 Hz) of the Leica SP8 confocal microscope was used to achieve a high spatiotemporal resolution. Frames were captured at a time interval of 5 to 15s. Optimal Z-size according to the objective was used (normally 1.19µm with pinhole airy1). Confocal stacks and movies were flattened by maximum projections using FIJI (ImageJ). The rest of the parameters remained the same as described in the confocal live-cell imaging section.

### Photoconversion (PhC)

Photoconvertible transgenic embryos from *Tg(neurod:kikume)* were kept as much as possible away from light to avoid spurious photoconversion, although basal levels are always present in our hands. Circular or hand-free designed ROIs the size of few cells was drawn in SP8 and SP5 inverted Leica microscope systems. In both microscopes, photoconversion was performed with UV 405 laser line at 5 to 15% diode power upon ROI at 200Hz during 6 scans and 3D ROI aiming the center of the photoconverted region to avoid same level of photoconversion from upper or lower planes, although inevitable to some extent. Bidirectional laser scanning on. Objective used Leica 506191 (20x) in glycerol position. PMT detectors employed. Phototoxicity controls were performed as follows: Laser power 100% at 200Hz during several scans (=>6). Cells do not seem to be affected, at least within 6-8 hours of recording.

### Photoablation (PhA)

SP5 inverted Leica Multiphoton was used. MAI TAI multiphoton activated (Mai Tai BB DeepSee [Spectra Physics] tunable [710–990 nm] pulsed laser). Humidity 4%, temperature 20°C. BS/RLD mirror and SP715 filter used. 910nm laser at 42% power, at 20x zoom with a HC PL APO 20x/0,75 immersion objective (#506191 from Leica) since no ROIs can be used in this confocal in multiphoton mode. PMT employed. Pinhole completely open (600nm). Scans ranged from 3 to 6 until a bubble formed (readout of destroyed tissue). Scan speed 200-400Hz. Frame size 1024×512, bidirectional ON, line and frame average 1.

### CRISPR Knock-in. Generation of *Tg(neurod:Gal4)* line (2 gRNAs)

crRNA sequences were selected using CHOP-CHOP (chopchop.cbu.uib.no). 2nmol crRNA were ordered to IDT (Integrated DNA Technologies) and 2-100nmol tracrRNA. Cas9 protein was ordered to IDT (Alt-R™ S.p. Cas9 Nuclease V3, 100 µg, #1081058). crRNA and tracrRNA were resuspended to a concentration of 100µM (20 µL) in IDTE buffer 1x (IDT, #11-05-01-14). 5 µL crRNA was mixed with 5 µL tracrRNA in 10 µL DUPLEX buffer IDT (IDT, #11-05-01-12), to get a final concentration of 25µM of duplex guideRNA (dgRNA). Mix is incubated for 5 minutes at 95°C and then cool down to room temperature. Aliquots might be stored at -20°C. The following mix was prepared: 0,86µL H20 RNAse free with 1µL of total dgRNA, 1µL of 250ng/ µL *gbait* sgRNA and 1,5 µL of 1ug/ µL Cas9. This mix is called RNP (Ribonucleoprotein complex). Warm RNP up at 37°C for 10 minutes. Add 0,5µL of the donor plasmid “gBait hsp70Gal4FF” (Kimura et al., 2014) at 200ng/µL. Final volume 5µL. The final conditions of such mix are: total dgRNA = 5,7µM = 200ng/µL; *gbait* sgRNA = 50ng/µL; Cas9 = 300ng/µL; donor plasmid = 20ng/µL. Followingly, inject 1nL per embryo.

References from the section: (Auer, Duroure, Cian, et al., 2014b; Auer, Duroure, Concordet, et al., 2014; Hoshijima et al., 2016, 2019; Kimura et al., 2014; Thomas & Raible, 2019).

### Genotyping fish from *Tg(neurod:Gal4)* line

*Tg(neurod:Gal4)* fish where either phenotyped crossing by UAS:Kaede or other UAS:reporter line, or by fin clipping and PCR genotyping with the following protocol: Anesthetized adult zebrafish (42µL tricaine per mL of water, from a stock of 400mg tricaine powder in 100mL H2O) were fin clipped. After that, genomic DNA extraction by using N-Amp kit extraction (XNAT2 Extract-N-Amp Tissue PCR Kit XNAT2-1KT). PCR protocol as follows: Fw primer Gal4FF: GCAGGCTGAAGAAGCTGAAG; Rv primer Gal4FF: GGAAGATCAGCAGGAACAGC. 94°C 3 minutes; 96°C 10 seconds; 57°C 15 seconds; 72°C 30 seconds; 72°C 10 minutes; hold at 4°C. 35x cycles. Product of 178bp.

### Tol2 injections (RhoGTPases, *Tg(neurod:kikume)* stable line generation)

Mix 1µL of Tol2 mRNA at 175ng/µL with 3µL of plasmid at 50ng/µL and 6µL of dH20 (Final volume =10µL). Inject 1nL into the cell. Injected quantity: 15pg of plasmid and 17,5pg of Tol2 mRNA. For transient expression experiments embryos were kept until experiment. For stable line generation, positive embryos were taken to the fish facility at 5dpf.

### CRISPR Eraser

To retain eGFP in just a few cells, we designed CRISPR Eraser. This methodology consists in <100% efficient Cas9 cutting and frameshift of eGFP reporter in a given transgenic line. To perform this protocol, mix 1,5µL of Cas9 at 1µg/µL (62µM) with 1µL of *gbait* sgRNA at 250ng/µL and 2,5µL of H2O RNAse free. Final volume 5 µL. Concentrations of sgRNA and Cas9 of 50ng/µL and 300ng/µL(6µM), respectively. Inject 1nL into 1 cell stage embryos into the cell or yolk (yolk injections will give more labeled cells). Injected quantity: 50pg sgRNA and 300pg Cas9. It is important that the transgenic line contains GFPbait sequence designed by (Kimura et al., 2014) for proper cutting. It can be done by sequencing or in vitro assay of a PCR product cutting. GFPbait sequence is GGCGAGGGCGATGCCACCTACGG.

### Quantifications

Nuclei were counted manually, measurements were done with FIJI (ImageJ V1.5) from a defined region of interest (ROI). When ROIs were used, the same area was used to compare samples. Cellular tracks were manually performed by using TrackMate (Tinevez et al., 2017). Directionality analysis and plot to origin tracks was performed using DiPER (Gorelik & Gautreau, 2014) according to (Breau et al., 2017).

### 2D Dispersion analysis

From merged-to-origin migratory plots (Fig. 3D, 4C and 6C’) obtained with DiPER (Gorelik & Gautreau, 2014), we extracted the relative spatial position in XY at endpoint of the recording of each NB. Final relative position in XY of each NB was plotted in RStudio (4.2.0) using the SDD function from the SIBER library obtained from (https://CRAN.R-project.org/package=SIBER) based on the pipeline in (https://cran.rproject.org/web/packages/SIBER/vignettes/Introduction-to-SIBER.html) (Jackson et al., 2011)(See Supplementary text1 for pipeline). SIBER library fits bi-variate ellipses to spatial data using Bayesian inference. Data of the ellipsoid accounting for the 95% confidence interval of data dispersion and ellipse centroid was retrieved and plotted with final relative position in XY of each NB in Excel.

### Kymographs

Onto 4D recordings (x, y, z, t) we drawn a line of 1 to 15 pixel thick in our region of interest, and followingly used the Kymograph option in FIJI to deploy a spatial section of an image with temporal resolution.

### Statistical analysis and plots

All the data in the work were first tested for normal distribution using the Kolmogorov-Smirnov test and the Levene’s test for homogeneity of variances. For two-group comparison two-tailed Student’s t test was used for parametric data or Mann-Whitney *U* test for non-parametric data. Values are expressed as mean ± SEM, mean ± SD or median. Graphs were done with GRAPHPAD PRISM 8 software. Error bars represent SEM or SD.

G*power3.1 (Erdfelder et al., 2009) program was used to infer *a priori* the sampling *n* needed to get statistically significant results from an expected phenotype of inferred penetrance.

## Acknowledgements

We thank members of the laboratory and the developmental biology group at UPF for insights and critical discussions (Laura Taberner, Nerea Montedeoca, Mireia Rumbo, Carolyn Engel-Pizcueta, Covadonga Fdez-Hevia) and technicians Laia Subirana and Marta Linares for technical support; the CRG-ALMU microscopy facility staff, PRBB zebrafish aquatics facility and UPF Genomics facility for technical support. We want to specially thank Dr. Esteban Hoijman for insights and corrections on this manuscript.

The authors would like to thank Dr. Katie Kindt for kindly providing Tg*(neurod:kikume)* construct; Dr. Verena Ruprecht and Dr. Esteban Hoijman for the *Tg(actb2:H2A- mCherry)* zebrafish line and the DN and CA RhoGTPases constructs (originally published in (Hanovice et al., 2016)); Dr. Cristina Pujades and Dr. Covadonga Fdez-Hevia for Gbait-Gal4 construct (originally published in (Kimura et al., 2014)); Dr. Jeroen Bakkers and Dr. Federico Tessadori for the UAS:H2A-GFP construct (Strate et al., 2015).

## Competing Interests

No competing interests declared

## Funding

This work has been supported by AEI-BFU2017-82723P, PID2020-117662GB-I00 (FEDER) from MCINN to BA and the Unidad de Excelencia María de Maeztu, AEI (CEX2018-000792-M). AB is a recipient of the predoctoral fellowship “Formación de Profesorado Universitario (FPU)” form the Spanish Ministry of Universities (FPU17/03287).

## Notes

### Competing Interest Statement

The authors have declared no competing interest.

